# DNA-binding-independent mechanisms of metabolic regulation by the *Drosophila* FOXO transcription factor

**DOI:** 10.1101/2025.08.20.670103

**Authors:** Victor Bustos, Lauren McDonagh, Bruna Digiacomo, Ralf L. Meilenbrock, Jacqueline Eßer, Sebastian Grönke, Cathy Slack, Linda Partridge

**Affiliations:** Max Planck Institute for Biology of Ageing, 50931, Cologne, Germany; Institute of Healthy Ageing and GEE, University College London, WC1E 6BT, UK; College of Health and Life Sciences, Aston University, Birmingham, B4 7ET, UK; School of Life Sciences, Warwick University, Coventry, CV4 7AL, UK

## Abstract

Forkhead box-O (FOXO) transcription factors are evolutionarily conserved regulators of several biological processes, including development, stress responses, metabolism and ageing. As downstream effectors of nutrient-dependent cell signalling pathways, including insulin/IGF signalling, they integrate signals from multiple stimuli to orchestrate appropriate transcriptional responses to changes in the nutritional environment. Traditionally, FOXO-dependent responses have been attributed to target gene regulation through direct interactions with regulatory regions by DNA-binding via the conserved Forkhead (FH) domain. However, emerging evidence suggests that FOXO proteins may also influence gene expression through DNA-binding-independent mechanisms. However, differences in transcriptional outputs between DNA-binding dependent and independent FOXO functions have yet to be explored. Here, we have used genomic engineering of the endogenous *Drosophila foxo* locus to disrupt the DNA-binding activity of the single fly FOXO orthologue, allowing us to dissect the *in vivo* contributions of canonical and non-canonical dFOXO functions. We show that while DNA-binding is essential for several dFOXO-mediated phenotypes including female fecundity, lifespan, and resistance to oxidative and xenobiotic stress, other traits such as adult body size and survival during starvation remain intact. Notably, DNA-binding-deficient dFOXO flies exhibit defective lipid mobilisation under starvation, implicating a DNA-binding-independent role for dFOXO in metabolic regulation. Differential gene expression analysis during starvation in these mutants revealed key transcriptional changes in genes encoding metabolic regulators as well as regulators of transcription and chromatin structure. Together, these findings reveal distinct modes of dFOXO transcriptional regulation that depend on its direct association with DNA.

## INTRODUCTION

The Forkhead box-O (FOXO) transcription factors represent a highly conserved family of transcriptional regulators that play key roles in a wide range of biological processes including cell proliferation, growth, metabolism, stress responses, reproduction and ageing (Cao *et al*., 2023; Santos *et al*., 2023; Rodriguez-Colman, Dansen and Burgering, 2024). All members of this family share the Forkhead (FH) DNA-binding domain with 90% sequence identity across orthologues, together with surrounding, largely unstructured N- and C-terminal regions. The FH domain consists of four alpha helices and three beta strands that adopt a winged helix structure. DNA binding is mediated via interactions between several specific amino acids within helix H3, for example amino acids N211 and H215 in human FOXOs, that interact with the same consensus DNA sequences in different species: the *C. elegans* DAF-16 family member binding element (DBE, 5′-GTAAA(T/C)AA-3′) and the insulin response element (IRE, 5′-(C/A)(A/C)AAA(C/T)AA-3′) (Obsil and Obsilova, 2011; Dai *et al*., 2021). The species-level conservation of FOXO DNA-binding is evident from *C. elegans* to mammals, as DAF-16 function is maintained when partially replaced by mammalian FOXO3a (Webb, Kundaje and Brunet, 2016).

As key downstream transcriptional effectors of reduced insulin/insulin-like growth factor (insulin/IGF) signalling, FOXO proteins are active under conditions of reduced nutrient input or starvation where they enter the nucleus to orchestrate an appropriate transcriptional response. Conversely, when nutrients are abundant and insulin/IGF signalling is active, FOXO proteins are negatively regulated by PI3K-AKT-dependent phosphorylation resulting in their nuclear exclusion and concomitant loss of transcriptional outputs. Many of the phenotypic responses to reduced insulin/IGF signalling in both *C. elegans* and *Drosophila* are dependent on DAF-16/dFOXO activation including lifespan extension, xenobiotic resistance and starvation survival (Yu and Larsen, 2001; Slack *et al*., 2011). This has led to an extensive search for direct DAF-16/FOXO target genes that are proposed to mediate at least in part some of the phenotypic effects of reduced insulin/IGF signalling (Alic *et al*., 2011; Kumar *et al*., 2015).

More recently, it has become apparent that FOXO proteins may also regulate gene expression through distinct mechanisms that do not rely upon their ability to bind to target genes directly (Kodani and Nakae, 2020). Initial experiments using human FOXO1 carrying a mutation within H3 of the DNA-binding domain (H215R) demonstrated that this DNA-binding deficient FOXO1 regulated a disparate set of genes compared to the unaltered FOXO1 protein (Ramaswamy *et al*., 2002). Subsequently, introduction of this same mutation into mice showed that murine FOXO1 controls glucose-related metabolism through its canonical transcription factor activity but alters lipid metabolism via both DNA-binding-dependent and -independent mechanisms (Cook *et al*., 2015). However, further explorations into the gene expression changes associated with these DNA-binding-independent roles in metabolism are still outstanding, alongside the mechanisms underlying these DNA-binding-independent functions and their possible presence in organisms other than mammals.

These non-canonical mechanisms of FOXO transcriptional regulation are most likely regulated through specific protein-protein interactions and several unrelated protein interaction partners for FOXOs have now been identified that together allow FOXOs to activate or repress the expression of a diverse array of target genes. These FOXO-protein interactions are proposed to mediate their transcriptional changes in a variety of ways including through chromosomal translocations, co-factor recruitment or sequestration, and co-factor displacement (van der Vos and Coffer, 2008). For many of these regulatory protein-protein interactions, FOXO is still required to bind directly to DNA. For example, FOXO1 interacts with the hepatocyte nuclear factor 4 (HNF-4), an essential factor in regulating carbohydrate metabolism, to repress expression of the glycolytic gene *GCK* and activate expression of the gluconeogenic gene *G6pc* but this still requires FOXO DNA-binding (Hirota *et al*., 2008). However, FOXO1 was found to displace the Notch signalling corepressors, nuclear corepressor (NcoR) and silencing mediator for retinoid and thyroid hormone receptor (Smrt) and recruit the coactivator mastermind-like 1 (Maml1) to initiate Csl activity and regulate the expression of cellular differentiation genes through interactions with Csl rather than direct regulation of target genes (Kitamura *et al*., 2007). Hence, FOXO-protein interactions can influence gene expression independently of direct FOXO DNA-binding, but how FOXOs use this mechanism to control other FOXO-dependent functions and the changes in gene expression these interactions cause remain relatively unknown.

Here, we have used genomic engineering to introduce amino acid substitutions within the *Drosophila* FOXO protein that disrupt its ability to bind to DNA and regulate direct target gene expression. Unlike in mammals, the *Drosophila* genome encodes a single FOXO protein which facilitates the *in vivo* study of FOXO function independently of redundancies between paralogues. We show that many of the physiological roles of *Drosophila* FOXO, including female fecundity, lifespan and survival under oxidative or xenobiotic stress require a functional DNA-binding domain. Interestingly, though, adult body size and starvation survival were normal in flies lacking functional FOXO DNA-binding activity. We demonstrate that FOXO-deficient flies under starvation have impaired mobilisation of lipid but not carbohydrate stores, a metabolic effect that occurs independently of dFOXO DNA-binding. Transcriptomic analysis of dFOXO-deficient versus dFOXO DNA-binding-deficient flies during starvation revealed overrepresentation of genes linked to processes associated with transcriptional regulation and chromatin structure in dFOXO DNA-binding-independent flies. Our results therefore describe metabolic functions of *Drosophila* FOXO that mediate normal starvation responses independently of its ability to directly bind to DNA.

## RESULTS

### Creation of DNA-binding deficient dFOXO

FOXO proteins across different species share a highly conserved DNA-binding domain (DBD) including two conserved residues that are in direct contact with DNA based on the crystal structure of FOXO1 (Brent, Anand and Marmorstein, 2008) (Figure 1A and 1B). To test whether these two amino acids in *Drosophila* dFOXO are important for DNA-binding and transcriptional activity, we expressed eGFP-tagged wild-type dFOXO (dFOXO-WT) or a DBD-mutated dFOXO protein in which both amino acids were changed to alanine (N146A and H150A) (dFOXO-DBD) in S2R+ cells. We first assessed the nuclear localisation of both dFOXO proteins in response to nutritional cues. Under nutrient starvation, both dFOXO-WT and dFOXO-DBD proteins were localised to the nucleus but promptly migrated to the cell cytoplasm upon stimulation with insulin (Figure 1C). We then evaluated the transcriptional activity of both proteins by co-transfection with two different reporter plasmids in which *luciferase* expression is under the control of two different promoters known to be bound by dFOXO: 4 x FOXO Responsive Elements (4xFRE) and the *Drosophila* Insulin Receptor (dInR) promoter. We observed significantly elevated luciferase expression from both reporter plasmids in cells expressing dFOXO-WT but not in cells expressing dFOXO-DBD (Figure 1D). Thus, mutation of these residues within the DBD of *Drosophila* dFOXO disrupts its transactivation activity but not its normal nuclear localisation.

**Figure 1.**
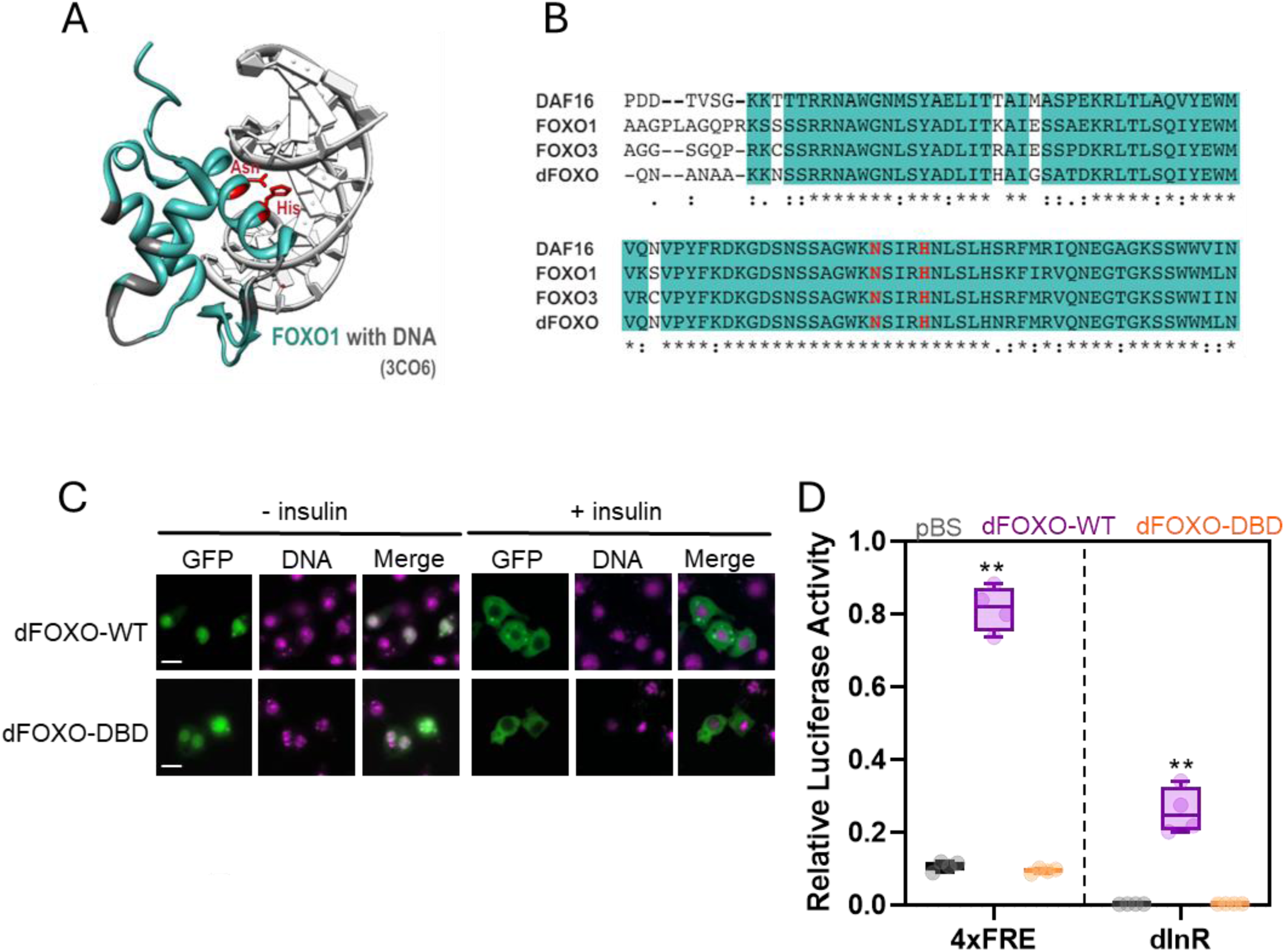
Mutation of key conserved residues that mediate FOXO-DNA interactions impair dFOXO transcriptional activity. (A) Crystal structure of FOXO1 DBD bound to DNA (PDB-ID: 3CO6) (Brent et al., 2008) visualized using the UCSF Chimera software (Pettersen et al., 2004). Blue residues are identical between FOXO1 and dFOXO. Red residues mediate the interaction between FOXO1 and DNA. (B) Protein sequence alignment of the DNA binding domains (DBD) of *C. elegans* FOXO (DAF16), *Drosophila* FOXO (dFOXO) and mouse FOXO1 and FOXO3. Identical amino acids shared between all organisms are shaded in blue. The two amino acids that are implicated in the interaction with DNA in FOXO1 are conserved across species and highlighted in red. (C) Representative fluorescent microscopy images of S2R+ cells transfected with GFP-tagged wild-type dFOXO (dFOXO-WT) or GFP-tagged dFOXO in which amino acids N146 and H150 are mutated to alanine to disrupt DNA binding (dFOXO-DBD). Transfected cells were maintained in serum-free media without (-) and with (+) insulin. Scale bar = 10 μm. (C) Luciferase reporter assays using S2-R+ cells containing either an insulin receptor (InR) promoter or an artificial promoter containing four FOXO responsive elements (4xFRE) upstream of luciferase. Cells were transfected with pBluescript (pBS) or expression plasmids for GFP-tagged dFOXO-WT or dFOXO-DBD. Box and whisker plots show the minimum and maximum value (bars), upper and lower quartiles (box), median (line within box). n = 4. ** *p*<0.05 (ANOVA with post-hoc Tukey’s multiple comparisons test).

### Genomic engineering of the *dfoxo* locus

To introduce the same DBD mutations as described above within the endogenous *dfoxo* locus, we first replaced ∼3kb of coding sequence, which includes most of the *dfoxo* ORF, by an attP-landing site using ends-out homologous recombination (Huang *et al*., 2009) to generate the *dfoxo^ΔV3^*parental line (Figure 2A). Phenotypically, this line was indistinguishable from the previously reported *dfoxo^Δ94^* null allele (Slack *et al*., 2011) when examined for developmental time, body weight, oxidative stress resistance and starvation survival (Supplementary Figure S1).

**Figure 2.**
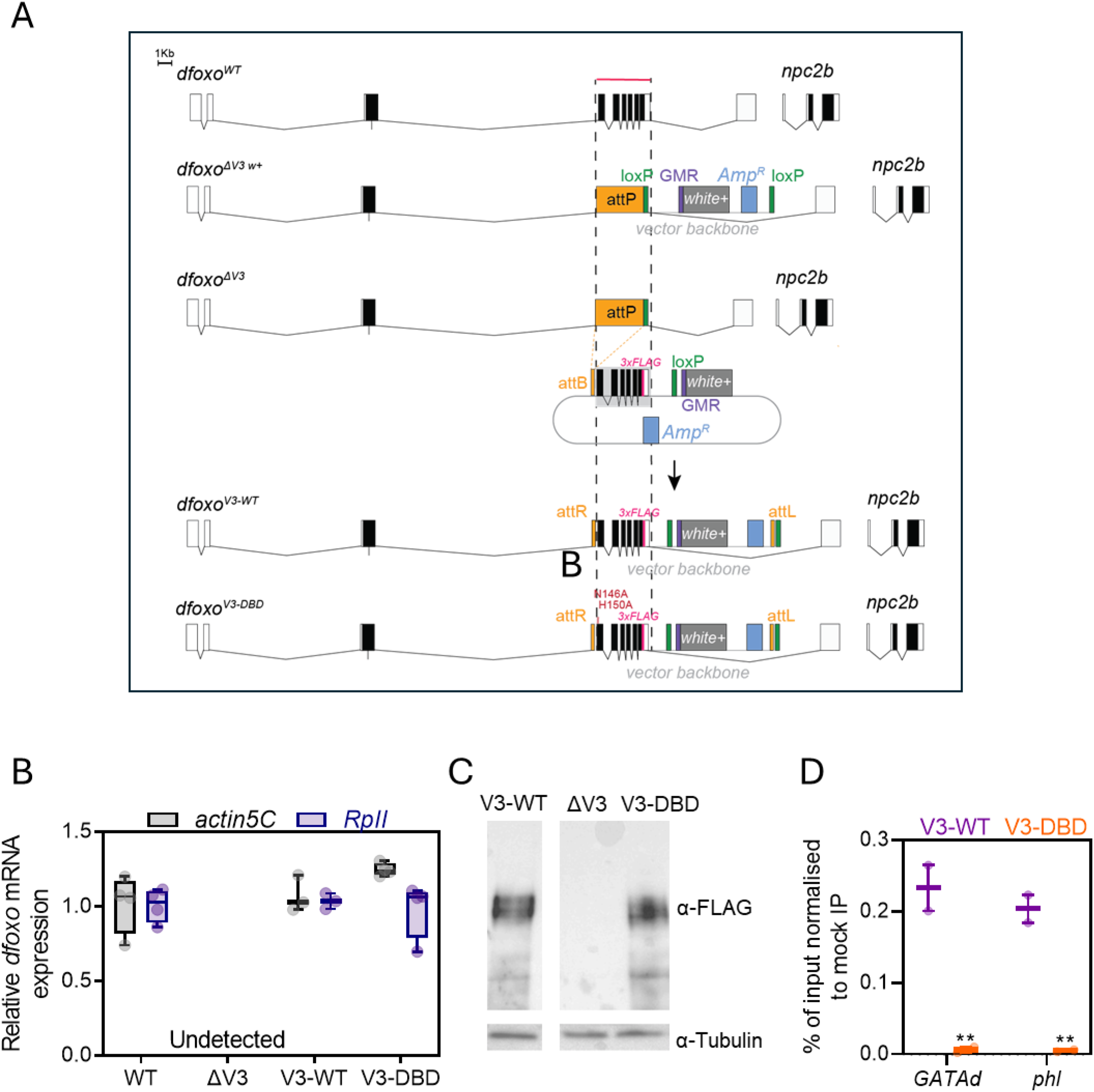
Genomic engineering of the *dfoxo* locus. (A) Schematic of the *dfoxo* gene locus and genomic engineering strategy to generate new knock-in alleles. Ends-out homologous recombination was used to replace the V3 (∼3Kb) region of the *dfoxo* locus (red line) with attP sites. After removal of the mini-*white* marker gene, the attP sites can be used to reintroduce modified versions of the *dfoxo* gene as shown here for the *dfoxo^V3-WT^* and *dfoxo^V3-DBD^* alleles. Coding (black) and non-coding (white) exons are represented by boxes and introns by lines. (B) qRT-PCR analysis of *dfoxo* mRNA expression in control (CTL), *dfoxo^ΔV3^* and the two reinsertion alleles, *dfoxo^V3-WT^* and *dfoxo^V3-DBD^* using probes against the *V3* region. *dfoxo* mRNA expression was normalised to *actin5C* or *RpII*. (C) Western blot on whole-body protein extracts from 10-day old female flies of the indicated *dfoxo* alleles probed with anti-FLAG and anti-tubulin antibodies. (D) ChIP-qPCR after immunoprecipitation using anti-FLAG antibody from extracts prepared from 7-day old adult females of the indicated genotypes. Data shows quantification of the abundance of DNA of known dFOXO target genes, *GATAd* and *phl,* associated with the immunoprecipitated material as a % of input chromatin isolated after ChIP normalised to a mock (no antibody) IP. (n = 2 chromatin extracts per genotype using 100 flies each, ** *p <* 0.05 (ANOVA with post-hoc Tukey’s multiple comparisons test)). Box and whisker plots show the minimum and maximum value (bars), upper and lower quartiles (box), median (line within box).

We then generated knock-in lines in which we introduced either a wild-type *dfoxo* sequence or a DBD-mutated sequence, both of which contained a carboxy-terminal FLAG-tag. All engineered lines were verified by sequencing. Expression of *dfoxo* mRNA in *dfoxo^V3-WT^* and *dfoxo^V3-DBD^* knock-in flies was comparable to control flies while *dfoxo* transcripts were undetectable within *dfoxo^ΔV3^*flies (Figure 2B). FLAG-tagged proteins of the corresponding size for dFOXO were detected at comparable levels in extracts from both the *dfoxo^V3-WT^* and *dfoxo^V3-DBD^* knock-in lines but not from *dfoxo^ΔV3^* flies (Figure 2C).

We then performed chromatin immunoprecipitation (ChIP) followed by qPCR targeted to the promoter regions of two dFOXO target genes, *GATAd* and *phl* (Alic *et al*., 2011). We found enrichment of the wild-type FLAG-tagged dFOXO protein at both promoters but not the DBD-mutated FLAG-tagged dFOXO (Figure 2D) suggesting that mutation of the dFOXO DBD prevents DNA binding *in vivo*.

### Phenotypic analysis of dFOXO DBD mutants

FOXO transcription factors are major effectors of insulin/IGF signalling and in both *C. elegans* and *Drosophila* are essential for many of the phenotypes associated with reduced insulin/IGF signalling (Kenyon *et al*., 1993; Larsen, 1993; Lin *et al*., 2001; Slack *et al*., 2011). To test whether any of these phenotypes also require a functional dFOXO DNA-binding domain, we examined the *dfoxo^V3-DBD^* mutants for their ability to restore any of the phenotypes associated with *dfoxo* loss-of-function. For adult lifespan, female fecundity, stress survival and feeding behaviour, the *dFOXO^V3-DBD^*mutants phenocopied the *dfoxo^ΔV3^* null flies (Figure 3A; Supplementary Figures S2 and S3). Note the phenotypes of *dfoxo^V3-WT^* flies were comparable to the background strain in these assays, demonstrating that the insertion of the FLAG tag does not affect dFOXO function. Interestingly, though, the *dfoxo^V3-DBD^* mutants did not show any obvious differences in adult body size and weight, and survival under starvation, unlike the *dfoxo^ΔV3^* null flies, which had reduced body size and weight and impaired survival under starvation (Figure 3B and C). Thus, these phenotypes are dependent on dFOXO activity but not on the presence of a functional DBD.

**Figure 3.**
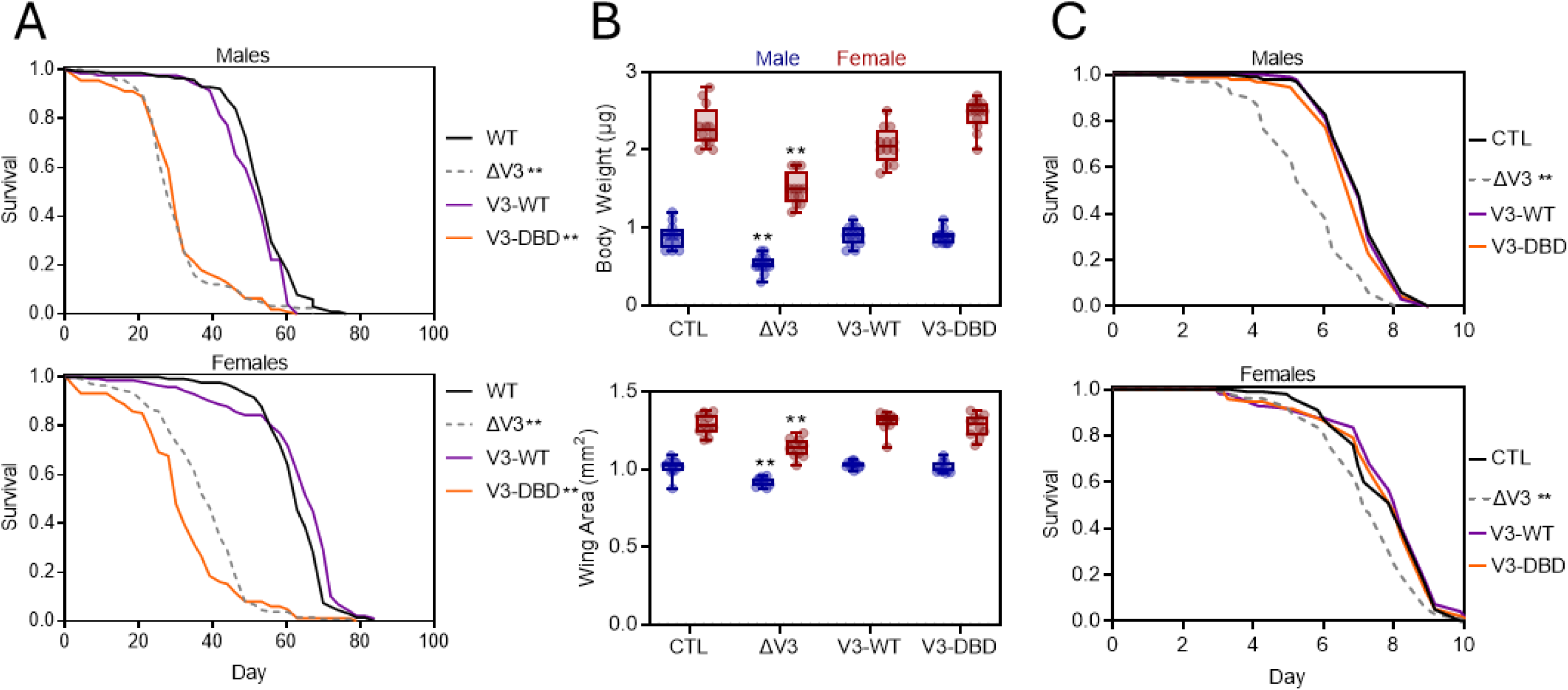
Requirements of the DNA-binding domain for *in vivo* functions of dFOXO. (A) Survival of control (CTL), *dfoxo^ΔV3^, dfoxo^V3-WT^* and *dfoxo^V3-DBD^* flies of the indicated sex. (n ∼ 100 flies per genotype). ** *p <* 0.001, Log-rank test vs *dfoxo^V3-WT^*. (B) Body weight and wing size measurements of 7-day old male and female flies of the indicated genotypes (n = 10 individual flies per genotype). Box and whisker plots show the minimum and maximum value (bars), upper and lower quartiles (box), median (line within box) with individual points overlaid. ** *p <* 0.05, ANOVA with post-hoc Tukey’s multiple comparisons test. (C) Starvation survival of control (CTL), *dfoxo^ΔV3^, dfoxo^V3-WT^* and *dfoxo^V3-DBD^* mutant flies of the indicated sex. (n ∼ 100 flies per genotype). ** *p* < 0.001, Log-rank test vs *dfoxo^V3-WT^*.

FOXO transcription factors are key regulators of both glucose and amino acid sensing (Housley *et al*., 2008; Green, Lamming and Fontana, 2022). The starvation sensitivity in *dfoxo^ΔV3^* null flies might therefore represent an inability to respond to nutrient deprivation at multiple levels. We therefore used a chemically defined diet in which we specifically removed either the sugar or amino acid components of the diet (Piper *et al*., 2014). In contrast to the *dfoxo^ΔV3^* null flies, which were sensitive to both sugar and amino acid starvation, the *dfoxo^V3-DBD^* mutants were sensitive to sugar but not amino acid starvation (Figure 4A and B). Together these data indicate that dFOXO regulates the response to sugar deprivation in a DNA-binding dependent manner, whereas it modulates the response to amino acid starvation independently of DNA-binding.

**Figure 4.**
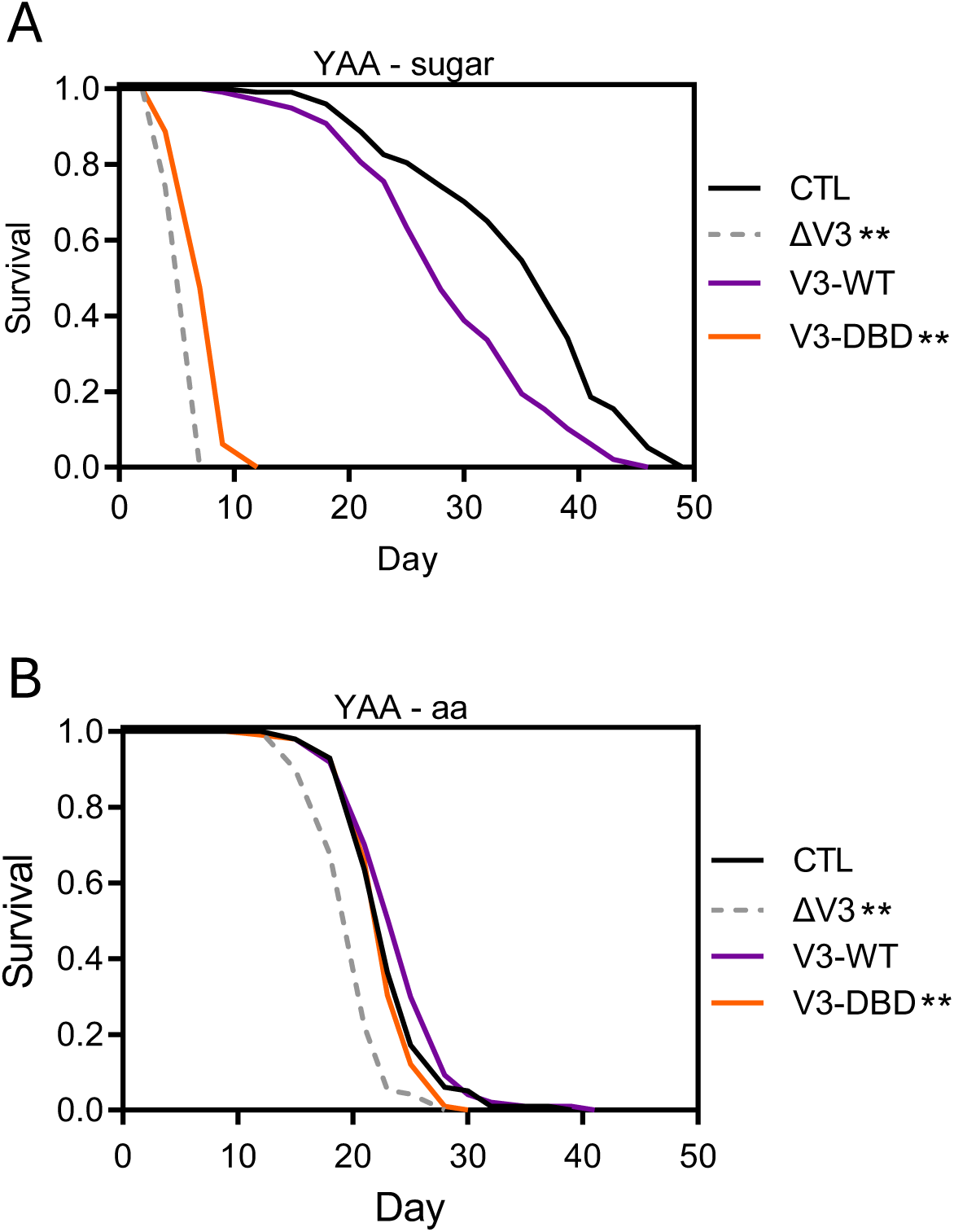
Sensitivity of *dfoxo* alleles to sugar versus amino acid starvation. Survival responses of control (CTL), *dfoxo^ΔV3^, dfoxo^V3-WT^*and *dfoxo^V3-DBD^* flies in the absence of either (A) sugar (YAA-sugar) or (B) amino acids (YAA-aa). n ∼ 100 flies per genotype. ** *p <* 0.001, Log-rank test vs *dfoxo^V3-WT^*.

### Changes in metabolic stores during starvation

Energy demanding processes, including growth during development and the ability to survive under nutrient deprivation, are particularly susceptible to metabolic dysfunction. This could manifest as differences in the ability to produce or maintain metabolic stores or to mobilise nutrient stores appropriately when required (Harshman and Schmid, 1998). *Drosophila* primarily store nutrients as trehalose, glycogen and triacylglycerides (TAG). We therefore performed quantitative measurements of these three metabolic storage molecules before and during starvation. No significant differences in the steady state stores of glycogen, trehalose or TAG were observed between the genotypes (Figure 5), suggesting that under normal nutritive conditions, both the *dfoxo^ΔV3^* null and *dfoxo^V3-DBD^* flies can produce and maintain normal metabolic stores. During starvation, trehalose levels did not significantly change for any of the genotypes (Figure 5A). In contrast, glycogen levels significantly decreased during the first 2 days of starvation, but with no significant differences between genotypes (Figure 5B). Interestingly, TAG levels were significantly reduced after 6 days of starvation for all genotypes except the *dfoxo^ΔV3^* null flies (Figure 5C). Thus, the reduced ability of *dfoxo^ΔV3^* null flies to survive under starvation may reflect an impaired ability to mobilise their TAG stores, which the *dfoxo^V3-DBD^*flies could do normally despite the absence of a functional dFOXO DBD.

**Figure 5.**
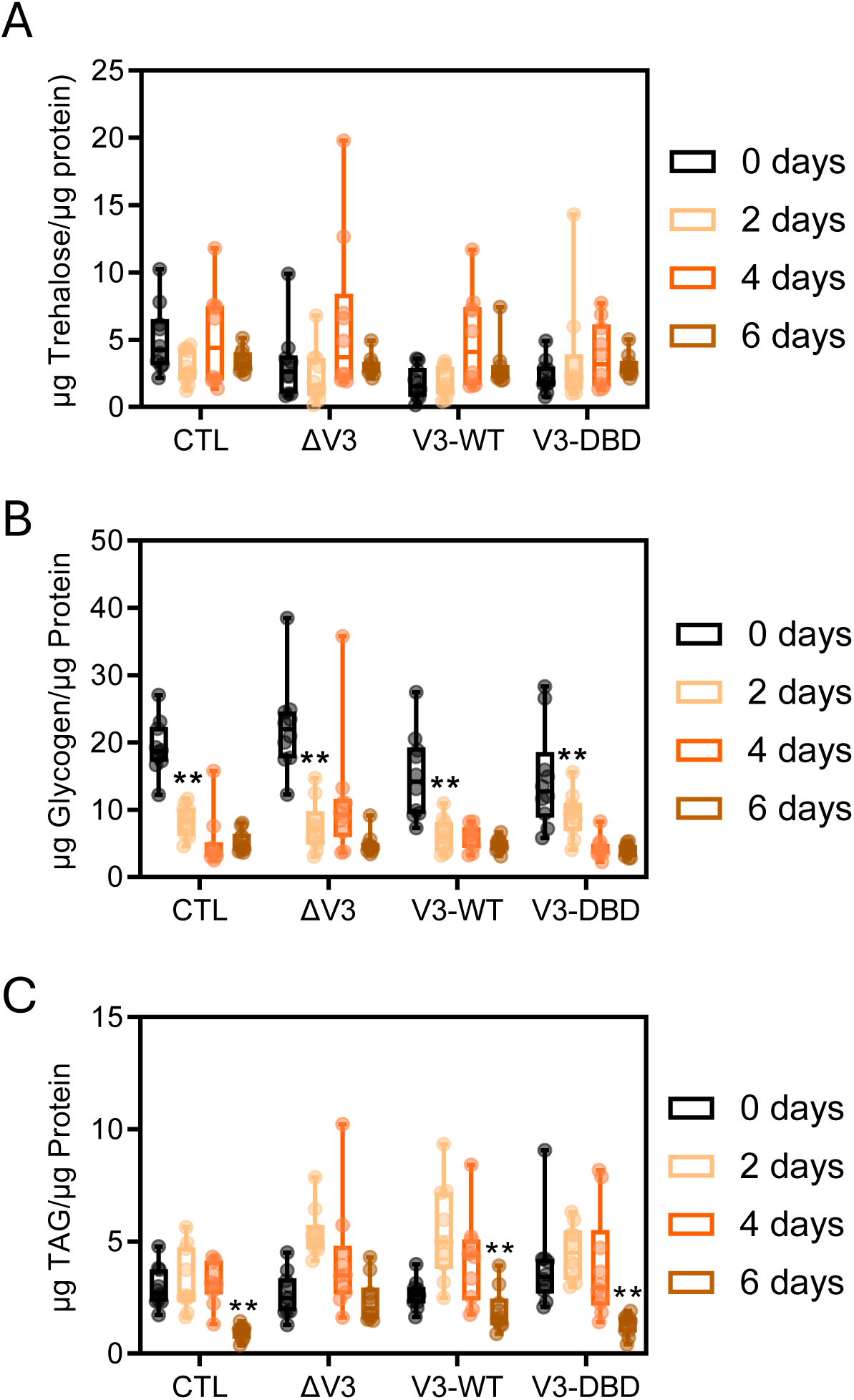
Lipid but not carbohydrate mobilisation during starvation requires dFOXO but not a functional DNA-binding domain. Extracts from adult female flies were assayed for (A) Trehalose, (B) Glycogen and (C) TAG content after 0, 2, 4, and 6 days of starvation with starvation commencing at 7-days of age. Concentrations of each metabolic storage molecule were normalised to protein content within the extracts. For all graphs, genotypes represented are control (CTL), *dfoxo^ΔV3^* (ΔV3), *dfoxo^V3-WT^* (V3-WT) and *dfoxo^V3-DBD^* (V3-DBD). Box and whisker plots show the minimum and maximum value (bars), upper and lower quartiles (box) and median (line within box). (n = 10 biological replicates per genotype of extracts from 5 pooled flies, ** *p* < 0.01, Generalised Linear Model with post-hoc Mood’s median test).

### Analysis of differential gene expression during starvation

To identify candidate mechanisms responsible for the differential survival responses of dFOXO-deficient versus DNA-binding deficient flies during starvation, we analysed starvation-induced genome-wide transcriptional changes in *dfoxo^ΔV3^, dfoxo^V3-WT^* and *dfoxo^V3-DBD^*flies using RNA-sequencing (Figure 6). Principal Component Analysis (PCA) demonstrated close alignment between replicate samples which clustered based on genotype and nutritional condition (Figure 6A). The expression differences between groups could be attributed to two principal components (PC1 and PC2) which together explained 72% of the total variation across data sets. A large number of genes were significantly differentially regulated (*p* < 0.05) upon starvation in each of the *dfoxo* genotypes (Figure 6B). We first validated the gene expression changes identified from our analysis by comparing to published data sets. We identified a significant overlap in the genes that were differentially expressed in *dfoxo^ΔV3^*vs *dfoxo^V3-WT^* flies to those previously identified as differentially expressed by microarray analysis in the *dfoxo^Δ94^* allele vs control flies (Alic *et al*., 2011) (Supplementary Figure S4A). Among those genes that were differentially expressed in the two *dfoxo* mutant alleles were several previously identified direct dFOXO target genes including the *Bigmax* transcription factor, the *Su(Hw)* chromatin insulator factor, several ribosome biogenesis genes including the *Tsr1* ribosome assembly factor, and the insulin/IGF signalling pathway component*, SOS* (Alic *et al*., 2011). Similarly, we found a significant overlap in the genes differentially expressed in *dFOXO^V3-WT^* flies after 72-hours of starvation with those identified in wild-type flies after 48-hours of starvation (Bauer *et al*., 2006) (Supplementary Figure S4B). Thus, the differentially expressed genes identified from our RNA-seq analysis correlate well with those described in other studies of similar genetic or nutritional manipulations.

**Figure 6.**
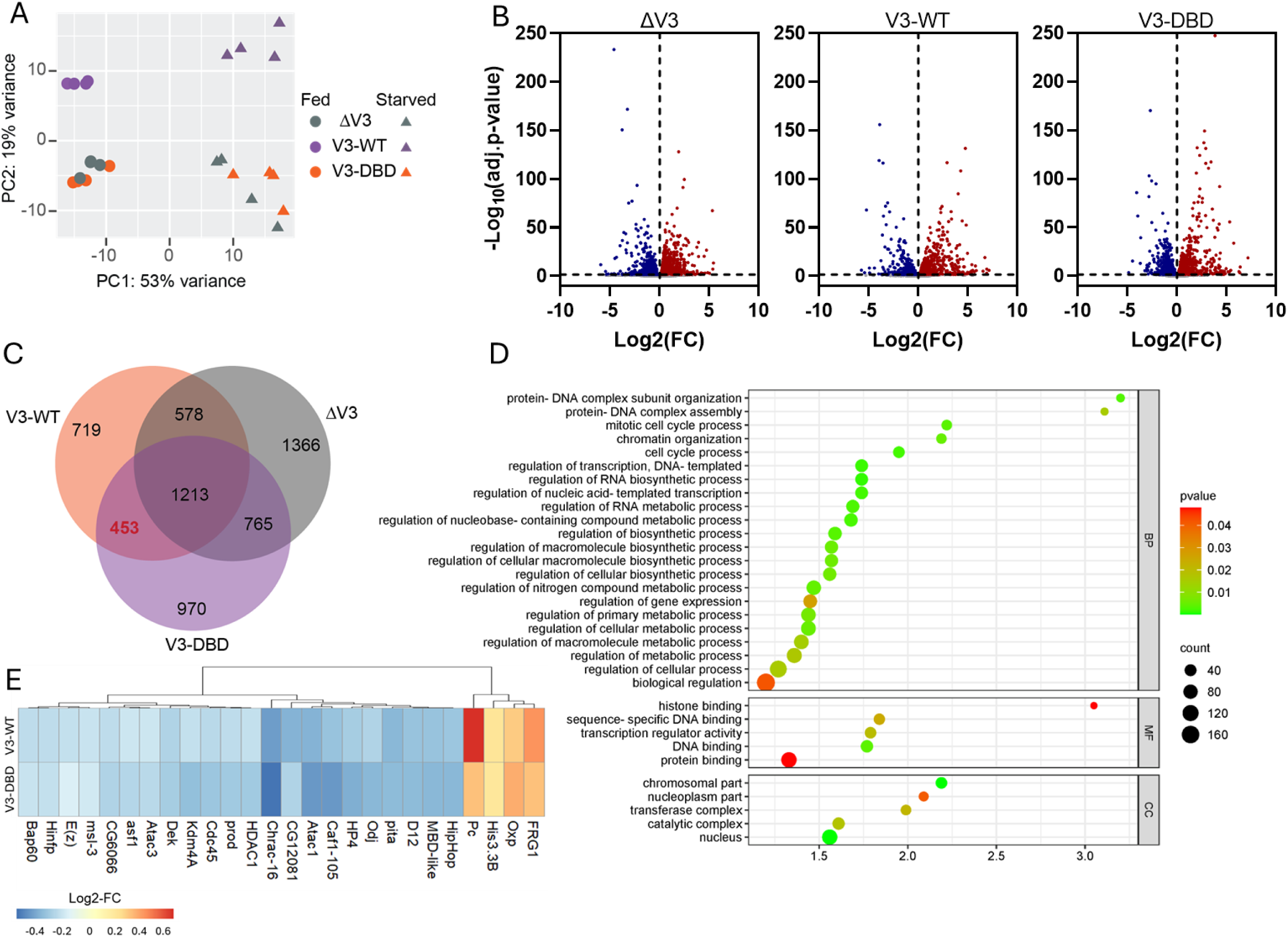
Starvation-induced gene expression changes in *dFOXO* mutant flies. (A) PCA of RNA-sequencing data from 10-day old adult females of the indicated genotypes when either fully fed or after 72 hours of starvation. *dfoxo^ΔV3^* (ΔV3), *dfoxo^V3-WT^* (V3-WT) and *dfoxo^V3-DBD^*(V3-DBD). (B) Volcano plots showing the number of differentially expressed genes in each *dfoxo* genotype during starvation (FDR 0.05%). Each dot represents an individual gene. Downregulated genes are shown in blue and upregulated genes are in red. Horizontal black dotted line denotes adjusted *p*-value threshold of –log_10_(0.05). (C) Proportional Venn diagram showing comparisons of the number of starvation-induced genes within the three *dfoxo* genotypes. Genes within the overlap regions are similarly regulated. 453 genes (red) were found to be differentially expressed during starvation in both *dfoxo^V3-WT^* and *dfoxo^V3-DBD^* flies but not in *dfoxo^ΔV3^* flies. (D) Gene Ontology (GO) enrichment analysis of the 453 dFOXO-dependent but DBD independent genes. Bubble plot shows the top GO terms (FDR 0.05%) of these overlap genes against a background list of all genes differentially expressed in both *dfoxo^V3-WT^* and *dfoxo^V3-DBD^*flies. (E) Heatmap of Log2FC for differentially expressed genes associated with chromatin organisation.

We then compared the genes that were differentially expressed after starvation in each of the *dfoxo* alleles. Approximately one third of these genes were shared across the three different *dfoxo* genotypes suggesting that a large proportion of the transcriptional response to starvation occurs independently of dFOXO (Figure 6C). Among these genes were several encoding regulators of both carbohydrate and lipid metabolism. For example, *Hexokinase-C*, *GFAT-2* and *Akh* which encode key factors involved in glycogen synthesis and storage were all differentially regulated after starvation in *dfoxo^ΔV3^, dfoxo^V3-WT^* and *dfoxo^V3-DBD^* flies. Similarly, all three *dfoxo* genotypes showed transcriptional changes within the expression of the TAG lipases *brummer (bmm)*, *doppelganger von brummer* (*dob*) and *Lip4* after 72-hours of starvation. Within the genes that were differentially expressed after starvation within the *dfoxo^V3-WT^* flies, ∼25% (719 genes) were not shared with *dfoxo^ΔV3^* or *dfoxo^V3-DBD^*flies indicating that the expression of these genes is modulated in a dFOXO-dependent manner during starvation (Figure 6C). Again, we identified several known direct dFOXO target genes in this group including the autophagy-related gene *ATG3* (Birnbaum *et al*., 2019) and the chromatin modifier *CTCF* (Alic *et al*., 2011).

To identify changes in gene expression mediated by dFOXO independently of its binding to DNA, we annotated the 453 genes that changed in expression during starvation in *dfoxo^V3-WT^* and *dfoxo^V3-DBD^* flies but not in *dfoxo^ΔV3^* flies (Figure 6C). Approximately 10% (47 of the 455) of the genes within this list were found to have unknown function or identity. Gene Ontology (GO) annotations enriched within the dFOXO-dependent but DBD-independent differentially expressed genes identified an overrepresentation of genes linked to multiple metabolic and biosynthetic processes as well as genes associated with chromatin modification and transcriptional regulation (Figure 6D). Among those annotated differentially expressed genes were several encoding UDP-glycosyltransferase enzymes (*Ugt37A1, Ugt37D1*, *Ugt37C2*, *Ugt316A1*, *Ugt36E1* and *Ugt49B1*) that catalyse the glycosylation of a broad range of lipophilic molecules including endogenous metabolites and signalling molecules to regulate their activity and distribution (Ahn and Marygold, 2021). Interestingly, expression of *Ugt35b* in glial cells has been recently shown to modulate lipid metabolism to influence survival in *Drosophila* (Sheng *et al*., 2025). Also among this group of differentially expressed genes were other key regulators of lipid and fatty acid metabolism including *mat, wal, DGAT2, scu, Kr-h1*, and *GPAT4*. Interestingly, several genes encoding known regulators of signalling downstream of the insulin receptor were within this group of differentially expressed genes including *gig*, *Sdr, Ptp61F* and *Lar*. dFOXO-mediated transcriptional feedback onto upstream intermediaries of the insulin signalling pathway has been previously reported (Puig *et al*., 2003; Alic *et al*., 2011; Slack *et al*., 2011) although our data suggests that this may not necessarily require direct association of dFOXO with the regulatory region of those target genes. We also identified several genes associated with chromatin modification and transcriptional regulation including the histone demethylase, *Kdm4*, and the histone deacetylase, *HDAC1*, which were both down-regulated in response to starvation in *dfoxo^V3-WT^* and *dfoxo^V3-DBD^* flies but unchanged in *dfoxo^ΔV3^* flies (Figure 6E) suggesting that dFOXO may modulate the transcriptional response to starvation through the modification of chromatin state to maintain survival independently of its ability to bind to DNA.

## DISCUSSION

FOXO transcription factors regulate multiple cellular processes by direct binding to target genes to regulate their expression. Using a novel genetic tool which abrogates *Drosophila* FOXO DNA-binding *in vivo*, we have shown that not all dFOXO-dependent regulation depends upon its ability to bind to DNA but that a subset of dFOXO functions, including regulation of adult body size and survival under starvation, do not require direct association of dFOXO with its target genes.

While these two processes are mechanistically distinct, they share a common requirement for extensive metabolic remodelling including the mobilisation of metabolic stores such as glycogen and TAG. Adult body size in *Drosophila* depends on the energy stores acquired during late larval development. These energy stores, primarily TAG stores, are subsequently used as metabolic fuel during metamorphosis (Merkey *et al*., 2011). Similarly, during chronic starvation adult *Drosophila* utilise their metabolic stores for survival, initially mobilising their carbohydrate stores and then switching to utilising TAG (Lee and Jang, 2014). Stored levels of both glycogen and TAG therefore reflect the metabolic balance of their biogenesis with catabolism. We observed no differences in either glycogen or TAG levels in fully fed dFOXO-deficient flies suggesting that dFOXO activity is dispensable for both glycogen and lipid synthesis under normal nutritive conditions. This was somewhat surprising as previous studies have shown that FOXOs modulate lipogenesis by disrupting transcriptional complexes to inhibit the actions of the sterol regulatory element-binding proteins 1c (SREBP1c), which are known to upregulate core lipogenic genes including *fatty acid synthase* (*FASN*) (Sekiya *et al*., 2007; Deng *et al*., 2012). However, other lipogenic pathways may be able to compensate for loss of dFOXO activity to regulate lipogenesis by converging on the same key regulatory genes. For example, the Mondo-Mlx signalling pathway, a key glucose-sensing pathway, can also induce lipogenesis and TAG storage in *Drosophila* through the activation of the transcription factor Sugarbabe, which in turn activates expression of both *FASN* and *AcetylCoA Carboxylase* (*ACC*), both key drivers of lipogenesis (Mattila *et al*., 2015). Interestingly, we found that *sugarbabe* expression was upregulated in fully fed *dfoxo^ΔV3^* versus *dfoxo^V3-WT^* flies in our RNA-seq analysis but no differences in the expression of *ACC* or *FASN* were observed in our transcriptomics data. Together, this supports the idea that multiple redundant mechanisms act in concert to regulate lipogenesis under normal dietary conditions.

In contrast, under starvation conditions, while initial carbohydrate usage was normal in the absence of dFOXO activity in *dfoxo^ΔV3^*flies, TAG mobilisation was delayed which may have contributed to the reduced survival of these flies during starvation. Starvation-induced TAG mobilisation appeared normal in FOXO DNA-binding mutants suggesting that these FOXO-dependent effects on TAG metabolism do not require direct binding of FOXO to target genes. In mammals, FOXO1 is thought to be a key promoter of the carbohydrate-to-lipid metabolic switch in skeletal muscle during fasting. When circulating glucose levels are low, FOXO1 transcriptional regulation facilitates a metabolic switch from carbohydrate oxidation to the utilisation of fatty acids for cellular energy through the upregulation of *pyruvate dehydrogenase kinase-4* (*PDK4*) and *lipoprotein lipase* expression (Cheng and White, 2011). Our results suggest that dFOXO mediates at least the initial phases of this metabolic switch in *Drosophila* but does so independently of a functional DNA-binding domain.

One potential mechanism through which dFOXO could mediate these effects independently of DNA binding is through modification to chromatin structure to modulate downstream gene expression. We identified enrichment of genes associated with histone binding and chromatin organisation within those genes differentially regulated during starvation in dFOXO DBD-deficient flies including key histone modifiers such as the histone lysine demethylase, *Kdm4*, and the histone deacetylase, *HDAC1*. The demethylase activity of Kdm4 is modulated in response to metabolic fluctuations as its key substrate, α-ketoglutarate, is a Kreb’s cycle intermediate (Tran, Lowman and Kong, 2017). This influence of metabolism allows Kdm4 to activate PDK4 indirectly via the E2F1 growth factor E2F1 leading to inactivation of pyruvate dehydrogenases and modulation of metabolic flexibility switching from glucose to fatty acid utilisation (Zhang *et al*., 2014). Overexpression of HDAC1 in human epithelial cells reduces lipogenesis through suppression of SREBP-1c promoter activity (Shin *et al*., 2021) while in *Drosophila* HDAC1 expression promotes starvation survival through increased expression of ribosomal RNA synthesis and autophagic genes (Nakajima *et al*., 2016). While the precise mechanisms through which dFOXO mediates its DNA-binding independent effects on cellular metabolism remain unresolved, differential expression of Kdm4 and HDAC1 would promote the switch from anabolic to catabolic metabolism as well as from carbohydrate to lipid utilisation during starvation and so may be central to the observed differences in starvation survival in dFOXO-deficient versus dFOXO DNA-binding deficient flies.

We have shown that flies fed a diet without sugar require dFOXO and its ability to bind to DNA to survive while in contrast survival on a diet deficient in amino acids requires dFOXO but not its canonical DNA-binding activity. FOXO transcriptional activity is directly regulated via nutritionally dependent post-translational modifications allowing for direct nutritional control of dFOXO-dependent gene expression. For example, AKT-dependent FOXO phosphorylation and inactivation is relieved under low glucose conditions (Ni *et al*., 2007) while the amino acid sensor GCN2 phosphorylates FOXO to promote its activity under conditions of amino acid deprivation (You *et al*., 2018). Furthermore, FOXOs contain multiple non-overlapping sites for posttranslational modifications other than phosphorylation including acetylation, glycosylation and ubiquitination, that regulate its activity in a combinatorial fashion (Zhao, Wang and Zhu, 2011). Posttranslational modifications to FOXOs have been shown to influence their interactions with other proteins. For example, FOXO3 association with the 14-3-3 chaperone proteins is dependent on its phosphorylation state (Singh *et al*., 2010). Increasing evidence suggests that FOXOs bind to a diverse array of different partner proteins including signalling molecules, other transcription factors and transcriptional cofactors (Kodani and Nakae, 2020). Differential posttranslational modification of FOXOs in response to changing nutritional cues may therefore influence their association with these partner proteins to affect downstream transcriptional regulation independently of direct FOXO DNA-binding. We have described here the transcriptional response to DNA-binding independent regulation of gene expression by dFOXO during starvation. Identifying the dFOXO protein partners that mediate these effects will provide a better understanding of the molecular networks through which these transcription factors integrate and interpret nutritional signals into complex physiological responses.

Our study therefore provides a powerful genetic tool for the *in vivo* manipulation of dFOXO DNA-binding activity in *Drosophila* and has revealed a new level of regulation for dFOXO during the starvation response based on its DNA binding capability. As metabolic regulation by FOXO factors is evolutionary conserved, it is likely that at least some of the potential FOXO interactions that modulate starvation in *Drosophila* will also be conserved in other species.

## EXPERIMENTAL PROCEDURES

### Drosophila stocks

The *dfoxo^ΔV3^* knockout founder line was generated by genomic engineering as previously reported (Huang *et al*., 2009; Kakanj *et al*., 2016). The *dfoxo* target region that we have named V3 (corresponding to 3R:14079252-14082284 of *Dmel* release r6.21) was substituted by a *white^hs^* marker gene and an attP-site using ends-out homologous recombination. ∼4Kb of flanking sequences of the target region were cloned into pBlueScript II SK (+) vector by ET recombineering (Zhang, Y *et al*., 1998; Muyrers *et al*., 1999) using a BAC clone that contains the *dfoxo* locus (CH321-24|13, BACPAC Resource Center, Oakland, California). After sequence verification, homologous arms were cloned into the pGX-attP targeting vector (Huang *et al*., 2009) and transgenic flies were generated by P-element mediated transformation using the BestGene *Drosophila* embryo injection service (Chino Hills, USA).

Crosses for ends-out homologous recombination were set for direct targeting as described before (Huang *et al*., 2008). The *white^hs^* marker gene was mapped to the third chromosome and homozygous flies were screened by PCR for the absence of the V3 region. We obtained one knockout founder (KO) line which was subsequently crossed with flies expressing cre-recombinase to remove the *white^hs^* marker gene (Groth *et al*., 2004). The resulting *w^[-]^* lines were crossed to flies expressing the ΦC31-integrase and used as the parental line for gene replacement at the *dfoxo* locus.

To generate *dfoxo* gene replacements, the V3 genomic region was cloned in the pBlueScript II SK (+) vector by ET recombineering and sequence verified. Reinsertion inserts were transferred to the pGEattBGMR vector (Huang *et al*., 2009) by restriction enzyme cloning using NheI and AscI to cut vector and insert. A vector containing the end of the last *dfoxo* coding exon and the beginning of the 3’UTR (3R:14081868-14082284, *Dmel* release r6.21) along with an in-frame sequence for a triple GS linker sequence and a 3xFLAG tag were generated by gene synthesis (Eurofins) and later In-Fusion cloned (Clontech) into the pBS-V3 vector that had been digested with BstEII and AscI. Mutations within the dFOXO DNA binding domain (DBD), were introduced in the pBS-V3-3xFLAG construct using QuickChange II XL site-directed mutagenesis (Agilent Technologies) and the V3-DBD-3xFLAG sequences were subsequently cloned in the pGEattBGMR vector. pGEattBGMR gene replacement constructs were then injected into ΦC31-integrase;*dfoxoΔV3w^[-]^* embryos at the transgenic fly core facility of the Max-Planck Institute for Biology of Ageing.

Other *Drosophila* stocks used in this study include *dfoxo^Δ94^*, a previously described *dfoxo* null allele (Slack *et al*., 2011). The *Drosophila* background strains, *white Dahomey* (*w^Dah^*) and *Wolbachia-*negative *white Dahomey* (*w^DahT^*) were used as control strains. *w^Dah^* was derived by incorporation of the *w^1118^* mutation into the outbred *Dahomey* background by backcrossing. The wild-type stock *Dahomey* was collected in 1970 in Dahomey (now Benin) and has since been maintained in large population cages with overlapping generations on a 12 h:12 h light to dark cycle at 25 °C. The *Wolbachia*-deficient *w^DahT^* strain was generated by treating *w^Dah^* flies with Tetracycline (25 mg/ml in standard SYA food) for three generations followed by a minimum of five generations to allow for full recovery from tetracycline treatment and restoration of intestinal flora. All mutations/transgenes were backcrossed for at least 6 generations into the corresponding background strain before experiments were conducted.

### *Drosophila* maintenance and husbandry

*Drosophila* stocks were maintained, and experiments were conducted at 25°C on a 12:12 h light:dark cycles at 65% humidity. Experimental flies were reared on standard sugar-yeast-agar food (SYA) (Bass *et al*., 2007) containing 5% (w/v) sucrose (Tate & Lyle), 10% (w/v) brewer’s yeast (MP Biomedicals) and 1.5% (w/v) agar (Merck) at a standardised larval density. Once emerged, adult flies were transferred to a fresh bottle containing SYA, allowed to mate for 24h and then sorted by sex under CO_2_ anaesthesia.

For lifespan assays, flies were sorted to a density of 15 flies per vial and transferred to fresh food 3 times per week while scoring for dead or censored flies. Flies were censored from the experiment if they were damaged during handling, escaped, or became stuck to the food. For fecundity measurements, 10-day old female flies were transferred to fresh SYA vials, removed 24 hours later and the number of eggs laid per vial were manually counted. Survival assays on YAA (defined *Drosophila* media) without sugar or amino acids were based on the 200N holidic medium as described in (Piper *et al*., 2014).

#### Stress Assays

Flies were maintained in SYA vials until 7 days post-eclosion before transfer to either 1% (w/v) agar in distilled water for starvation assays, 5% (v/v) hydrogen peroxide (in 1.5% (w/v) agar, 5% (w/v) sucrose) or 20 mM paraquat (in SYA media) for oxidative stress assays, or 0.03% (w/v) DDT (in SYA media). Deaths were recorded every 4 hours, 4 times per day and flies were transferred to fresh media every 2-3 days.

#### Developmental timing and body size measurements

Protocols for the measurement of developmental timing and body size were as essentially described before (Grönke *et al*., 2010). For developmental timing, eggs were collected over a 3h period and transferred to SYA food at a density of 50 eggs per vial. As the adult flies eclosed, the number of flies emerging was counted at regular intervals. To measure body weight, Individual adult flies were immobilised by incubation on ice and weighed using a BM-5D analytical balance. The left-hand wing was then removed from each individual fly and mounted in 50% (v/v) glycerol/ethanol. Wings were imaged using a Leica MZ10F stereo microscope coupled to a QImaging QIclick Digital CCD Camera using Micro-Manager 1.4 software. Wing areas were measured from the digitised images using the polygon tool in ImageJ between standardised points.

#### Capillary feeder (CAFÉ) assay

Capillary feeder (CAFÉ) assays were conducted as previously described (Ja *et al*., 2007). Individual 7-day old flies were transferred to a 7 mL bijou containing approximately 1 mL of 1% (w/v) agar. Graduated glass capillary tubes were filled with liquid fly food media (consisting of 5% (w/v) sucrose, 2% (w/v) autolysed yeast and Brilliant Blue dye. The tubes were then sealed with mineral oil to prevent evaporation and the capillary inserted into the top of the bijoux. The level of food within the capillary was then measured every 24 hours for a total of 96 hours.

#### Trehalose, Glycogen and TAG assays

Trehalose and glycogen concentrations were measured from extracts of whole 7-day old adult flies. Samples of 5 adult flies were homogenised in 50 μL 0.2 M Na_2_CO_3_ and incubated at 95°C for 2 hours before the addition of 30 μL 1 M acetic acid and 120 μL 0.2 M Na acetate (pH 5.2). For trehalose measurements, 4.2 units of trehalase (Megazyme) were added before incubation at 37°C overnight. For glycogen measurements, 0.0096 units of amyloglucosidase from *Aspergillus niger* (Merck) were added before incubation at 57°C overnight. Trehalose and glycogen standards of known concentration were similarly processed in parallel. Liberated glucose was then measured using the Infinity Glucose reagent (ThermoFisher Scientific) according to the manufacturer’s instructions.

TAG concentrations were also measured from extracts of whole 7-day old adult flies as described previously (Grönke *et al*., 2007). Briefly, five flies per sample were homogenised in 0.05% (v/v) Tween20 in PBS. The homogenate was then incubated at 70°C for 5 minutes. Glycerol standards of known concentration were processed in parallel. 5 µL of standard or sample were then incubated with 200µl of Infinity Triglycerides reagent (ThermoScientific) as per the manufacturer’s instructions.

Protein concentrations were also measured for each sample using the Pierce 660nm Protein Assay reagent.

#### Quantitative real-time PCR (qRT-PCR) analysis

Total RNA from 7-day-old adult flies was extracted using standard Trizol (Invitrogen) protocols. 1 µg of total RNA was reverse transcribed using SuperScript VILO (Invitrogen) followed by DNAse treatment (Ambion). Quantitative real-time PCR was performed using Taqman probes (Applied Biosystems) in a 7900HT real-time PCR machine (Applied Biosystems). Relative gene expression was determined by the ΔΔCT method and normalized to *actin* (*act5C*) or *RpII*. Four independent biological replicates per genotype were analyzed.

#### Chromatin immunoprecipitation (ChIP) and qPCR

Chromatin extraction and immunoprecipitation was performed as previously described (Alic *et al*., 2011). For chromatin extraction, ∼200 7-day-old flies were homogenized in PBS supplemented with protease inhibitors and 0.5% formaldehyde. Samples were quenched by the addition of glycine. Chromatin was collected by centrifugation and washed twice with FA/SDS buffer (50mM Hepes-KOH pH7.5, 150mM NaCl, 1mM EDTA, 0.1% (v/v) Na Deoxycholate, 0.1% (v/v) SDS, 1% (v/v) Triton-X 100 and freshly added 1mM PMSF). Chromatin was sheared by sonification to an average size ∼300bp and immunoprecipitations were performed using mouse anti-FLAG M2 primary antibody (Sigma, #F3165) and Dynaneads (Invitrogen). Mock IPs were performed in parallel in which no antibody was added to the sample. Immunoprecipitated chromatin was eluted from the Dynabeads using Pronase enzyme (Sigma). Samples were then treated with RNAse A (Qiagen) and the chromatin purified using the Qiagen PCR purification columns. Chromatin was then used as a template for quantitative PCR. Target genes were chosen based on (Alic *et al*., 2011).

## *Drosophila* cell culture

*Drosophila* S2R+ cells were cultured on Schneider’s Medium (Gibco) supplemented with 10% FCS, penicillin and streptomycin. Transfections was performed using Qiagen-Effectene® transfection reagent following the manufacturer’s instructions. The *dfoxo* ORF was cloned into the pUbiP-EGFP-rfA plasmid (kindly donated by Dr. Alf Herzig) using the Gateway system (ThermoFisher). Mutations within the *dfoxo* CDS were introduced using QuickChange II XL site-directed mutagenesis (Agilent Technologies). Cells were co-transfected with the pACT-renilla plasmid (obtained from Dr. Michael Hoch) and either the pGL-InR or pGL-4xFRE (4xFOXO Responsive Element) luciferase plasmids (as reported in (Puig *et al*., 2003). Measurements of firefly and renilla luciferase were performed 24h after transfection using the Promega Dual-Glo assay. For cell imaging, Cells were grown in a µ-slide (Ibidi) and starved on serum-free media for 2 hours prior to imaging using a fluorescent microscope (Leica). After the initial imaging, cells were treated with 10 µg/ml insulin for 10 min and imaged again. Hoechst 33342 was used to stain the nuclei.

### Western blot analysis

5 whole adult flies were lysed in 100 µL Laemmli buffer containing 100 mM DTT (1,4-Dithiothreitol) and proteins were denatured at 85 °C for 10 min. ∼20 μg of proteins were separated using 10% SDS–PAGE gels and transferred onto a 0.45-μm nitrocellulose membranes. Primary antibodies used were anti-FLAG (diluted 1:1,000; Sigma #F3165) and anti-tubulin (diluted 1:5,000; Abcam #ab108342). Horseradish peroxidase-conjugated secondary antibodies – either goat anti-rabbit IgG (1:10,000 dilution; Abcam #ab6721) or goat anti-mouse IgG (1:10,000 dilution; Abcam #ab6789) were used depending on the corresponding species of the primary. Blots were developed using the Clarity ECL detection system (Bio-Rad). Signals were captured using a G:BOX gel documentation system (SynGene) and band intensity was quantified using Fiji (Schindelin et al., 2012).

### RNA Sequencing

Total RNA was extracted from whole 10-day old adult female flies using TRIzol as described before. Flies were either fully fed or had been starved for 72 hours. cDNA libraries were prepared and sequencing (150 bp paired-end) performed by Novogene Co. Ltd (UK) using Illumina NovaSeq platforms. Sequences were quality checked by FastQC and reads below the minimum quality threshold Q<10 were discarded. Orphaned reads after filtering were also discarded. Reads were mapped to the Berkeley *Drosophila* Genome Project (BDGP) *Drosophila melanogaster* genome assembly release 6 (Ensembl release v51) using RNA STAR (Dobin *et al*., 2013). Reads that were uniquely aligned to annotated genes were counted with FeatureCounts version 1.6.2. Differential expression analysis was performed using DESeq2 version 1.10.1. Significance was determined using the standard DESeq2 FDR cutoff of adjusted *P <* 0.05. Gene overlaps were visualised using proportional Venn diagrams in BioVenn (Hulsen, de Vlieg and Alkema, 2008) and analysed using hypergeometric analysis in Excel based on the modified Fisher Exact p-value test, Expression Analysis Systematic Explorer (EASE). Gene ontology (GO) analysis was performed using GOrilla (Eden *et al*., 2009).

### Statistical analysis

Sample sizes were determined based on previous publications using similar experimental strategies. No statistical method was used to predetermine sample size. No specific methods were used to randomly allocate samples to groups. Data distributions were assumed to be normal, but this was not formally tested. No data were excluded from the analysis. Data collection and analysis were carried out in an unblinded fashion. Statistical analyses were mainly performed in R studio (R v3.5.5), Prism (v10.0, Graphpad) or Jmp (v14, SAS). Survival data were analysed using Log-rank test in Excel (Microsoft). Other statistical tests used are indicated in the figure legends. Error bars indicate SEM. Box-and-whiskers plots, show min to max whiskers.

## DATA AVAILABILITY

RNA-seq data files are openly available in ArrayExpress (accession number E-MTAB-15493). All other data that support the findings of this study are available from the corresponding author(s) upon reasonable request.

## ACKNOWLEDGEMENTS

V.B was funded through a Boehringer Ingelheim Fonds (BIF) PhD fellowship and L.M was funded by an Aston University 50th Anniversary PhD studentship. This project also received funding from the Max Planck Society to LP. We would like to thank Dr. Alf Herzig, Dr. Gábor Júhasz, Dr. Michael Hoch and Dr. Nazif Alic for providing reagents as well as all members of the Partridge and Slack labs for useful discussions. We acknowledge the Bloomington *Drosophila* Stock Center, the Vienna *Drosophila* Resource Center (VRDC) and FlyORF for providing fly strains, and the BACPAC Resources Center for providing BAC clones.

## AUTHOR CONTRIBUTIONS

V.B, S.G, and L.P designed the study. L.M and C.S designed and performed the RNA-sequencing experiments. J.E. performed embryo injections to generate some of the transgenic fly lines. V.B, L.M, B.D and R.M performed the experiments. V.B and L.M analyzed the data. V.B, S.G, C.S and L.P wrote the manuscript.

---

**Supplementary Figure S1.**
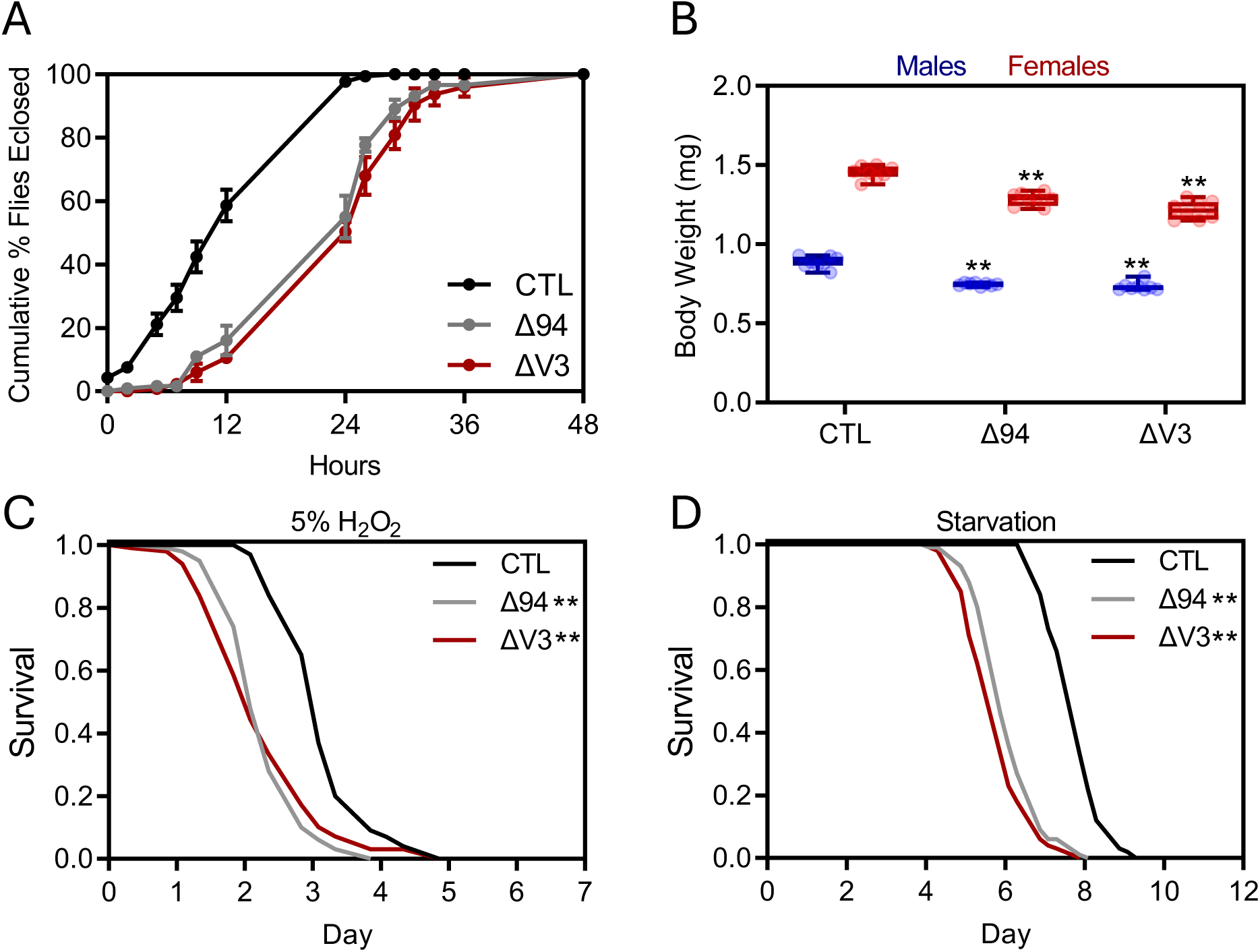
Phenotypic comparison of *dfoxo^ΔV3^ flies* to the *dfoxo^Δ94^* null allele. (A) Cumulative % of flies eclosing as adults from vials of the indicated genotypes (n=50 larvae per vial, 4 vials per genotype). Data indicates means ± SEM. (B) Body weight measurements of 7-day old male and female flies of the indicated genotypes (n = 8 individual flies per genotype). Box and whisker plots show the minimum and maximum value (bars), upper and lower quartiles (box), mean (line within box) with individual points overlaid. ** *p <* 0.05, ANOVA with post-hoc Dunnet’s multiple comparisons test. (C) Survival of 7-day old female flies of the indicated genotypes in the presence of 5% (v/v) hydrogen peroxide. (n = 99 to 100 flies per genotype. ** *p* < 0.0001, Log rank test compared to control (CTL) flies. (D) Survival of 7-day old female flies of the indicated genotypes during starvation. (n = 99 to 100 flies per genotype. ** *p* < 0.0001, Log-rank test compared to control (CTL) flies. For all graphs, genotypes represented are control (CTL), *dfoxo^Δ94^* (Δ94) and *dfoxo^ΔV3^* (ΔV3).

**Supplementary Figure S2.**
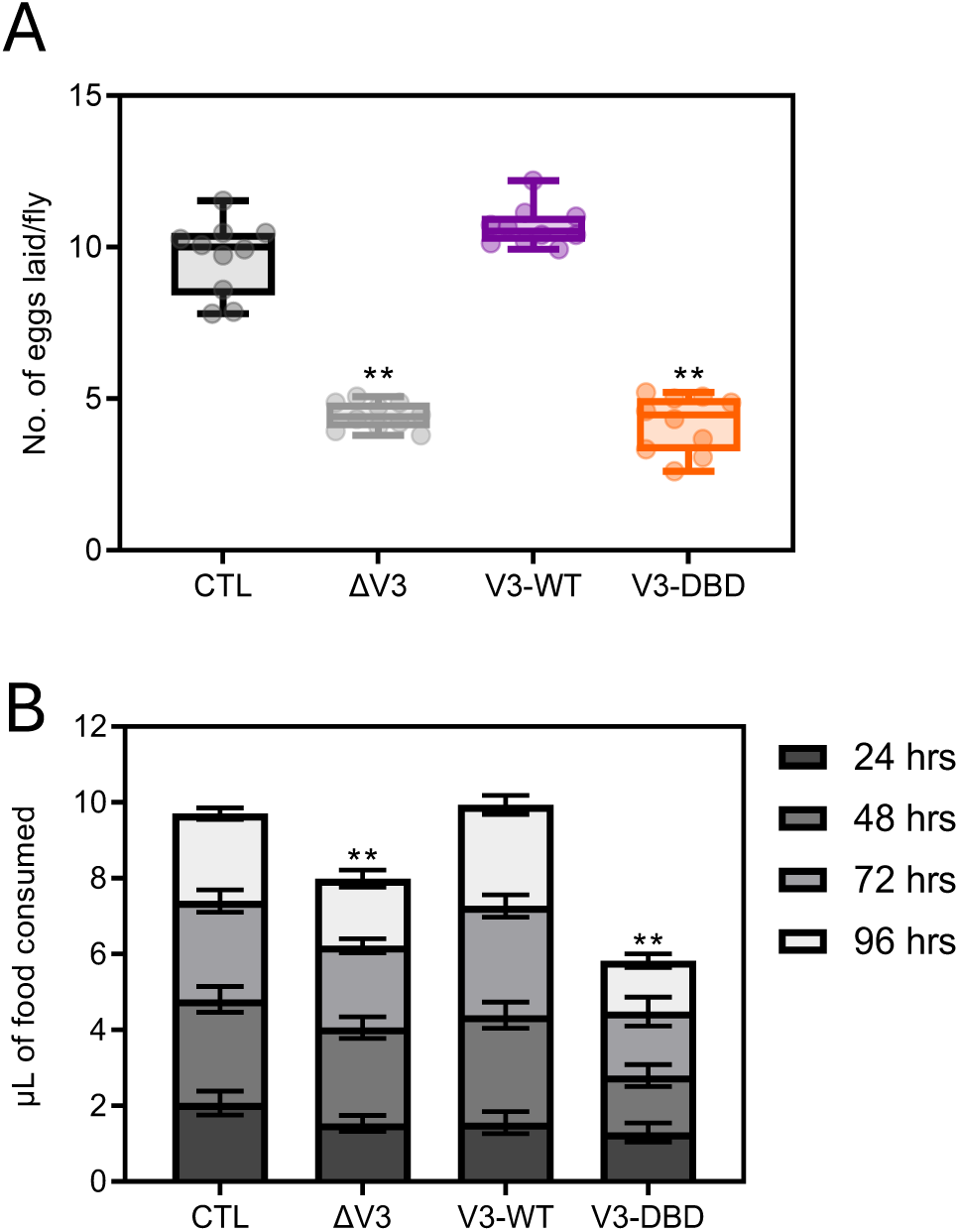
Normal fecundity and feeding require a functional dFOXO DNA-binding domain. (A) Number of eggs laid per fly over 24-hours for 10-day old control (CTL), *dfoxo^ΔV3^, dfoxo^V3-WT^* and *dfoxo^V3-DBD^* females. Box and whisker plots show the minimum and maximum value (bars), upper and lower quartiles (box), median (line within box) with individual points overlaid. ** *p <* 0.05, ANOVA with post-hoc Tukey’s multiple comparisons test. (B) Volume (in µL) of food consumed by individual flies of the indicated genotypes measured using the CAFÉ assay at 24, 48, 72 and 96 hours (n = 10 individual flies per genotype). Stacked bar graphs show mean values ±SEM. ** *p <* 0.05, generalised linear model with post-hoc Mood’s median test.

**Supplementary Figure S3.**
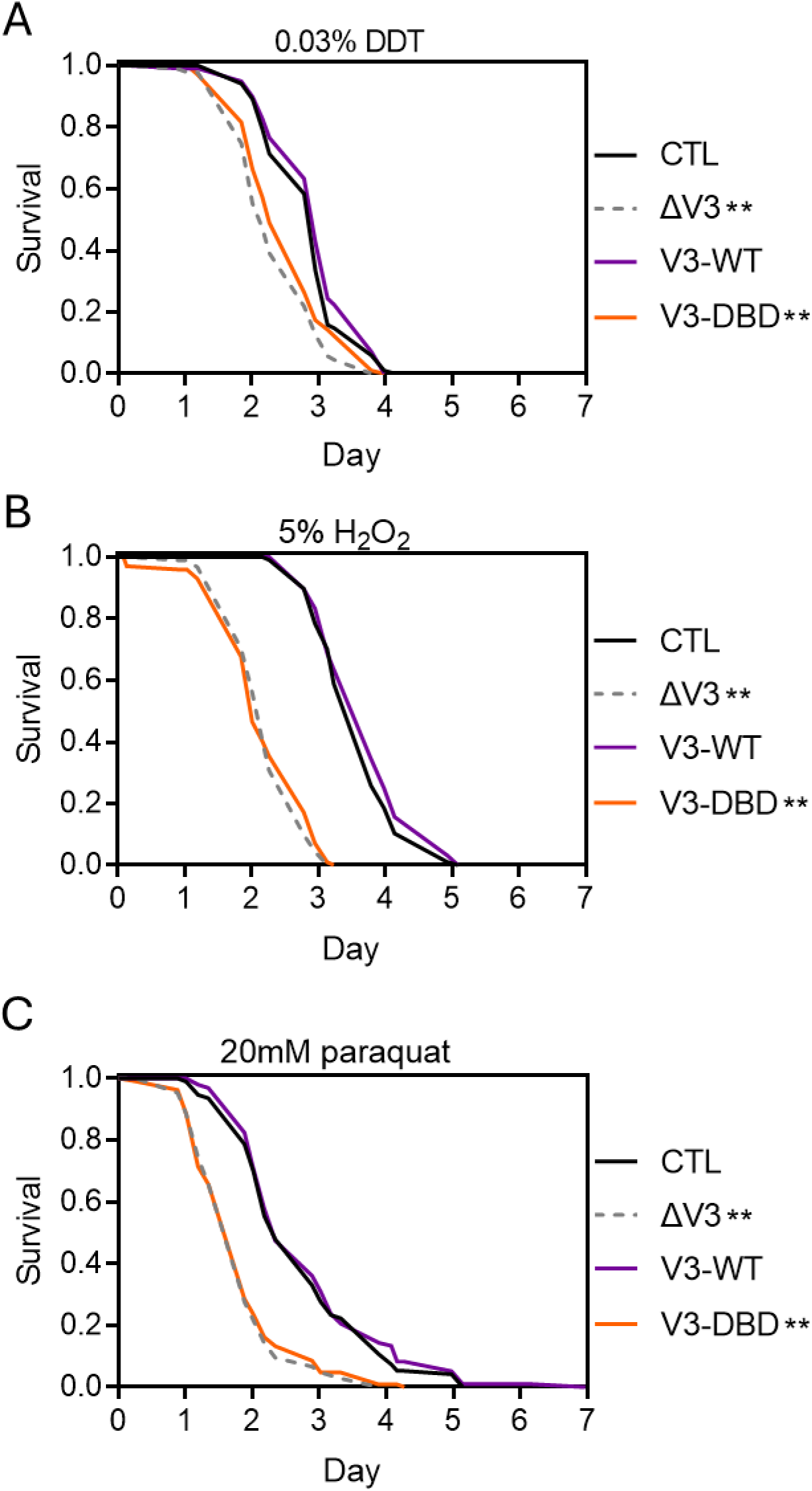
dFOXO DNA-binding domain is required for normal responses to xenobiotic and oxidative stresses. (A) Survival of 7-day old female flies of the indicated genotypes in the presence of 0.03% (w/v) DDT (n = 80 to 100 flies per genotype. ** *p* < 0.0001 **, Log-rank test compared to control (CTL) flies). (B) Survival of 7-day old female flies of the indicated genotypes in the presence of 5% (v/v) hydrogen peroxide. (n = 95 to 100 flies per genotype. ** *p* < 0.0001 **, Log rank test compared to CTL flies). (C) Survival of 7-day old female flies of the indicated genotypes in the presence of 20mM paraquat. (n = 94 – 108 flies per genotype. ** *p* < 0.0001 **, Log rank test compared to CTL flies). For all graphs, genotypes represented are control (CTL), *dfoxo^ΔV3^* (ΔV3), *dfoxo^V3-WT^* (V3-WT) and *dfoxo^V3-DBD^* (V3-DBD).

**Supplementary Figure S4.**
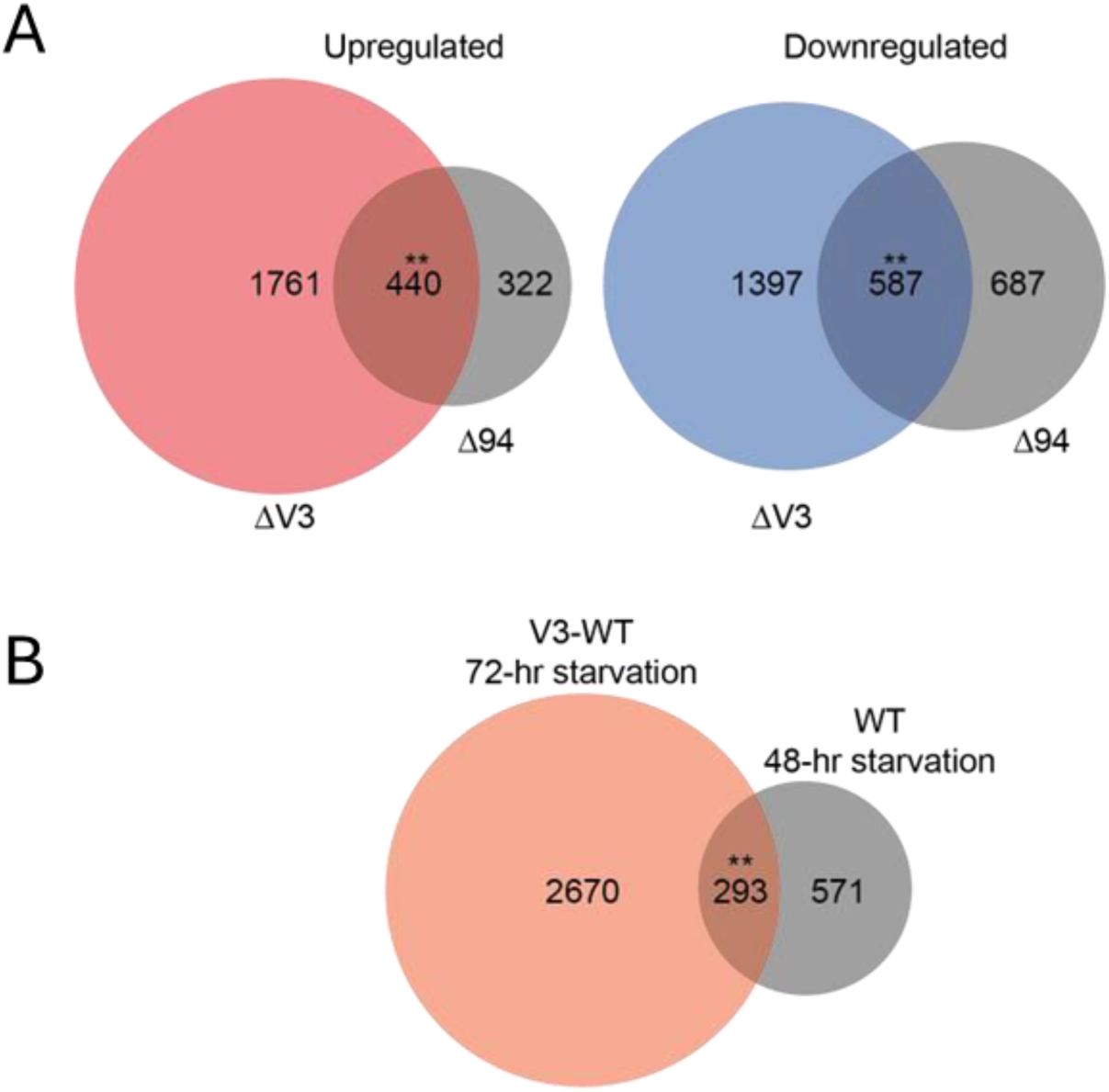
Differential gene expression analysis of *dfoxo* alleles. (A) Proportional Venn diagram comparing the differentially expressed genes in fully fed *dfoxo^ΔV3^* versus *dfoxo^V3-WT^* flies to those identified in *dfoxo^Δ94^* mutants (Alic *et al.,* 2011). Significant overlaps were identified in both the up- and down-regulated genes (** up-regulated genes, *p* = 1.38 x 10^-162^; down-regulated genes, *p* = 1.22 x 10^-182^, EASE analysis). Numbers indicate the number of genes in each category. (B) Proportional Venn diagram comparing the differentially expressed genes in *dfoxo^V3-WT^*flies after 72-hours of starvation to those of wild-type flies after 48-hours of starvation as described in (Bauer *et al.,* 2006). (** *p* = 1.40 x 10^-17^, EASE analysis). Numbers indicate the number of genes in each category.

